# Activity of natural occurring entomopathogenic fungi on nymphal and adult stage of *Philaenus spumarius*

**DOI:** 10.1101/2023.07.14.548874

**Authors:** N. Bodino, R. Barbera, N. Gonzalez-Mas, S. Demichelis, D. Bosco, P. Dolci

## Abstract

The spittlebug *Philaenus spumarius* (Hemiptera: Aphrophoridae) is the predominant vector of *Xylella fastidiosa* (Xanthomonadales: Xanthomonadaceae) in Apulia and Europe. Current control strategies of the insect vector rely on mechanical management of nymphal stages and insecticide application against adult populations. Entomopathogenic fungi (EPF) are biological control agents naturally attacking spittlebugs and may effectively reduce population levels of host species. Different experimental trials in controlled conditions have been performed to i) identify naturally occurring EPF on *P. spumarius* in Northwestern Italy, and ii) evaluate the potential for biocontrol of the isolated strains on both nymphal and adult stages of the spittlebug. Four EPF species were isolated from dead *P. spumarius* cadavers collected in semi-field conditions: *Beauveria bassiana, Conidiobolus coronatus, Fusarium equiseti* and *Lecanicillium aphanocladii*. All the fungal isolates showed entomopathogenic potential against nymphal stages of *P. spumarius* (≈ 45 % mortality), except for *F. equiseti*, in preliminary trials. No induced mortality was observed on the adult stage. *Lecanicillium aphanocladii* was the most promising fungus and its pathogenicity against spittlebug nymphs was further tested in different formulations (conidia vs blastospores) and with natural adjuvants. Blastospore formulation was the most effective in killing nymphal instars and reducing the emergence rate of *P. spumarius* adults, reaching mortality levels (90%) similar to those of the commercial product Naturalis®, while no or adverse effect of natural adjuvants was recorded. The encouraging results of this study pave the way for testing EPF isolates against *P. spumarius* in field conditions and find new environmentally friendly control strategies against insect vectors of *X. fastidiosa*.

## 1. INTRODUCTION

The spittlebug *Philaenus spumarius* (L.) (Hemiptera: Aphrophoridae) is the key vector of the phytopathogenic bacterium *Xylella fastidiosa* Wells et al. (Gammaproteobacteria: Lysobacteraceae) (*Xf*) in Europe (Cornara et al. 2018, 2019; Cavalieri et al. 2019). *Xf* is a xylem-limited bacterium identified as the causal agent of several plant diseases in the American continent (e.g. Pierce’s disease on grapevine and citrus variegated chlorosis on citrus), and its recent discovery in the Mediterranean basin has led to serious phytosanitary concerns. *Xf* was first identified in Europe in 2013 in Apulia (South Italy), and the strain *X. fastidiosa* subsp. *Pauca* ST53 was characterized as the causal agent of the olive quick decline syndrome (OQDS), a severe plant disease that led to a dramatic dieback of olive trees in the Italian Region (Cariddi et al. 2014; Almeida 2016; Saponari et al. 2017). Several *Xf* strains have been then found in different Mediterranean countries (Italy, Spain, France, Portugal, Israel, Lebanon) – although with less severe economic impact than Apulia’s scenario – as a results of separate introduction routes of different subspecies and strains, some dating back to forty/fifty years ago (Soubeyrand et al. 2018; Moralejo et al. 2020; Dupas et al. 2023). The economic and social consequences of *Xf* in Apulia have already been significant (Saponari et al. 2019; Italia Olivicola Consorzio Nazionale 2019) and they could increase exponentially in the near future, given that the bacterium continues spreading in the Region (Kottelenberg et al. 2021). In addition, the pathogen and its associated diseases could be introduced to other olive-producing areas of the Mediterranean or produce new pathosystems with other susceptible crops, e.g. grapevine (Sicard et al. 2018; Landa et al. 2020; Schneider et al. 2020, 2021).

*Xf* is transmitted from plant to plant by xylem-sap feeding insects, and *P. spumarius* is by far the prominent vector in Europe (Almeida et al. 2005; Bodino et al. 2019; Cornara et al. 2019). Thus, the spittlebug represents a primary target of control strategies to reduce the spread and impact of *Xf* epidemics, given the current lack of curative applications upon persistent systemic infections of crop plants (Morelli et al. 2021; Bodino et al. 2023).

The control of spittlebug populations in Apulia relies on mechanical treatments, such as tillage and mowing, to kill nymphs in spring, and application of insecticides on olive trees against the adult stage (EFSA Panel on Plant Health (PLH) et al. 2019; Sanna et al. 2021; Morelli et al. 2021; EU 2020/1201). While the effectiveness of agronomic measures in reducing spittlebug populations in the field is demonstrated (EFSA Panel on Plant Health (PLH) et al. 2019; Sanna et al. 2021; López-Mercadal et al. 2022), the impact of insecticidal treatments on vector density remains unclear. Indeed, several issues with the use of insecticides against spittlebugs are currently present, among them: i) few active ingredients available, also due to the recent banning of some neonicotinoids, the most effective chemical family of insecticide against *P. spumarius* (Dongiovanni et al. 2018), ii) quite low persistence of available insecticides, such as organophosphates and Spinosad (Dáder et al. 2019), iii) high densities of both nymphal and adult stage in semi-natural reservoirs (Bodino et al. 2019; Cornara et al. 2021; Cappellari et al. 2022); iv) adult spittlebugs exposed to insecticides could present sublethal effects and/or active movements (flight), possibly impairing the effectiveness of control measures to on disruption of *Xf* transmission (Bodino et al. 2020; Lago et al. 2022). Moreover, insecticidal treatments may have not only a significant economic impact but also harmful consequences on non-target arthropods, on microbial biodiversity (Coll et al., 2011; Schreck et al., 2012), deep and surface water (Bony et al., 2008), animal and human health (Tsakirakis et al., 2014; Raherison et al. 2019).

Alternative biological control strategies against Xf vectors are currently investigated and their implementation strongly suggested by EU in the framework of the European “Green Deal”, that requires a transition to fair, healthy and environmentally friendly agricultural systems (https://ec.europa.eu/info/strategy/priorities-2019-2024/european-green-deal_it). The adverse environmental effects of chemical insecticides have increased the interest in microbiological agents over the years. Among others, entomopathogenic fungi (EPF) stand out as efficient biological control agents that infect the insect hosts by contact through the cuticle. Most, if not all, insect species can be naturally infected by epf, which exploit them as a nutrient source (Lovett and St. Leger 2017). As natural pathogens of insect pests, fungi have been successfully used as biocontrol agents in various crops (Rondot and Reineke, 2020) and about 70% of organic products are mycopesticides (Vidal and Jaber, 2015). They are mainly based on fungal propagules to be applied on the vegetation through nebulizing treatments which ensure almost complete coverage and maximize the probability of contact and infection of the target insect.

EPF have been found associated in field conditions to some American *Xf* vectors, e.g. the glassy-winged sharpshooter *Homalodisca vitripennis* (Germar) and *Oncometopia facialis* (Signoret) (Hemiptera: Cicadellidae: Cicadellinae). Various fungi species are known to infect these insects with various efficiency, e.g. *Hirsutella homalodiscae* nom. Prov. (Ascomycota: Hypocreales), *Beauveria bassiana* Balsamo Vuill. (Ascomycota: Hypocreales), *Isaria poprawskii* Caban., J.H. de Leon, Humber, K.D. Murray & W.A. Jones (Ascomycota: Hypocreales), *Metarhizium anisopliae* (Metschn.) Sorokīn (Ascomycota: Hypocreales)*, Trichothecium roseum* (Pers.) Link (Ascomycota: Hypocreales), although some are probably secondary pathogens or saprobes rather than primary pathogens of the sharpshooter (Kanga et al. 2004; Boucias et al. 2007; Pria Júnior et al. 2008; Dara et al. 2008; Lietze et al. 2011; Cabanillas and Jones 2013).

Scientific literature is rather poor in information on EPF natural occurring on *P. spumarius*. In the past, Weaver and King (1954) reported that an epidemic of *Entomophthora aphrophorae* Rostr. was observed in Denmark by Sorauer in 1913. Also, *Zoophthora petchi* was found attacking *P. spumarius* in England (Ben-ze’ev and Kenneth, 1981) and Holland (Mietkiewski and Van der Geest, 1985) but no further evidence has been reported in literature to date. Only recently, EPF strains isolated from environment and belonging to *Trichoderma* genus have been successfully tested for their pathogenicity against *P. spumarius* (Ganassi 2022), although under laboratory conditions.

In light of the above, a gap of knowledge on EPF species naturally occurring on *P. spumarius* needs to be filled. The possibility of handling native EPF strains isolated directly from the target insect could represent an improvement in terms of both awareness and success rate, and strains isolated from indigenous cadavers warrant study of their biological control potential (Shang et al. 2022, Quesada-Moraga et al. 2023).

Thus, the aims of this study were i) the identification of naturally occurring EPF isolated from *P. spumarius* cadavers in semi-field conditions and ii) the evaluation of biocontrol potential of the strains showing promising pathogenic attitude against both nymphal and adult stage of the target insect *P. spumarius*.

## 2. MATERIALS and METHODS

The present study span over 4 years, from 2019 to 2022, because of both the univoltine life cycle of the target insect and the COVID-19 pandemic restrictions in 2020. In 2019, *P. spumarius* individuals affected by mycosis were recovered, and fungal strains isolated and identified. In late spring-early summer 2019, preliminary tests were carried out using isolates belonging to the most promising fungal species, against both *P. spumarius* nymphs and adults. Trials were resumed in 2021, after COVID-19–related interruption, and focused on the pathogenic activity of *Lecanicillium aphanocladii* strain versus *P. spumarius* nymphs. Finally, the trials were replicated in spring 2022.

### 2.1 Origin and identification of EPF strains

#### 2.1.1 Fungi from infected P. spumarius adults

Fungal strains were isolated from *P. spumarius* adults affected by mycosis and recovered from a semi-field maintenance rearing in Torino. Seven *P. spumarius* cadavers found externally covered by mycelium were carefully collected and each placed in a micro tube with a brush sterilized with alcohol. The subsequent handling of insects and fungi was performed in laboratory under aseptic conditions: portions of mycelium emerging from cadavers were transferred, by using lancet and needle, to Sabouraud dextrose agar (SDA, Oxoid, Milan, Italy) Petri plates, supplemented with penicillin G (0.1 mg/mL) and streptomycin (0.5 mg/mL) prepared in sterile distilled water and added after autoclaving the medium. SDA 90-mm plates were incubated at 25 °C and mycelium growth checked every 2 days; fungal isolates were subcultured and pure cultures were obtained by single spore isolation (monosporic cultures).

Fungi were, then, observed under an Olympus BH-2 microscope with phase contrast at 400× magnification and three out of seven were discarded as they showed the characteristic somatic and multiplicative structures of the saprophytic genus *Rhizopus* sp. The four remaining fungal strains were transferred to mycological tubes on Malt Extract Agar (MA, Oxoid) slants, and kept at 4 °C for their storage.

#### 2.1.2 Molecular identification of fungi

The four fungal strains stored at 4 °C were transferred to MA Petri plates and cultured at 25 °C for 10 days. For each strain, 100 mg of mycelium was harvested and transferred in a micro tube for DNA extraction. Fungal genomic DNA was extracted using DNeasy Plant mini kit (Qiagen, Milan, Italy) in accordance with the manufacturer’s instructions. The internal transcribed spacer (ITS) region was used to identify the fungi as proposed by Schoch et al. (2012). PCR was performed by using ITS1 and ITS4 primers (White et al., 1990) and amplicons were sent for purification and sequencing to Eurofins GmbH (Hamburg, Germany). The generated sequences were then analysed using NCBI BLAST database (GenBank) to find the closest match of the sequences for fungal species identification.

### 2.2 Collection and rearing of *P. spumarius*

Early nymphal stages (1^st^-2^nd^) of *P. spumarius* were collected directly in the field during the early Spring (March) of each year of the study. Samplings were carried out on herbaceous cover in a mixed agricultural area near Torino (Chieri: 45.0179 N, 7.787 E). Nymphs were gently collected from the ground vegetation and placed into mesh/plastic containers with excised *Taraxacum officinalis* L. leaves for transportation to the laboratory facilities. Upon arrival at the laboratory, nymphs were transferred to potted *Cichorium intybus* L. (six true leaves growth stage) and *Vicia faba* L. (four true leaves) plants in mesh/plastic cages (Bugdorm BD4S4590) located under controlled conditions in climatic chamber (21 °C and 14:10 h L:D photoperiod). Nymphs were maintained in mass rearings under controlled conditions until they reached the appropriate developmental stage to start the fungal pathogenicity bioassays.

### 2.3 Screening of EPF strains in preliminary trials on *P. spumarius* nymphs and adults

#### 2.3.1 Fungal strains and preparation of conidia inoculum

The four fungal strains previously isolated and identified as entomopathogenic species were used in preliminary trials performed in 2019 to assess their potential pathogenic activity against *P. spumarius* nymphs and adults. For each strain, a conidia inoculum to be used in the tests was prepared as follows: the fungal strains were transferred from MA slants kept at 4 °C to Potato Dextrose Agar (PDA, Oxoid) Petri plates and incubated at 25 °C for 14 days; then, conidia were collected with a spatula by scraping them from the surface of mycelium flooded with 2 mL of sterile distilled water, and placed in sterile tubes. The suspensions were then filtered through sterile gauze to remove mycelial debris and, after that, conidia concentrations were evaluated at optical microscope by counting at Bürker chamber. Suspensions were adjusted with sterile distilled water to obtain the final volume of 15 mL (5 mL per replica, for each strain) and the concentration of 1*10^8^ conidia/mL, as reported by other authors for controlled condition trials (González-Mas et al. 2019, Moussa et al. 2021). The suspensions were added with 0.05 v/v Tween 80® (v/v) to break the surface tension of water and evenly distribute the conidia, and vortex-mixed for 2–3 min.

The suspensions were freshly prepared on the day of the trials when also conidia viability was determined based on germ tube formation (Cabanillas et al. 2013). Briefly, 10 μL of properly diluted conidial suspensions were spread, with a sterile spatula, on a thin layer of PDA (Oxoid) in 45 mm Petri plates. Plates were incubated at 25 °C and conidia germination assessed at transmitted light microscope after 24 h. Subsamples of 100 conidia were scored and they were deemed to have germinated when the longitude of the germ tube was longer than half the size of the conidia. Conidia suspensions were judged acceptable if the viability was 90% and above.

#### 2.3.2 Experimental setup

The entomopathogenic activity of the four fungal strains previously isolated was tested in 2019 on *P. spumarius* nymphs and adults by direct spraying of conidia suspensions. Nymphal and adult stages were exposed to five experimental theses corresponding to the four fungal strains and a negative control (water plus 0.05 v/v Tween 80® (v/v), *Wtr*). Experimental microcosms (i.e., replicas) consisted in two *V. faba* plants (3 true leaf-age) sown in a single rounded pot (diam. 14 cm) enclosed by a clear plastic cylindrical cage (diam. 14 cm, height 45 cm, two holes with mesh for ventilation at top and side). The cylindrical cage was specifically designed to tightly fit inside the pot to ensure no space allowing insect escape.

Ten *P. spumarius* individuals (2^nd^-3^rd^ stage nymphs or adults emerged no more than 30 days before, according to the assay) were introduced in each microcosm, after being randomly collected from mass rearings. Three replicas per thesis were performed. Each microcosm was nebulized by aerograph with 5 mL of suspension (1 × 10^8^ conidia/mL = 5 × 10^8^ conidia in total) to reach the runoff point on plant leaves. The fungal treatments were applied 24 h after placing the nymphs on plants, in order to allow nymphs to settle on leaves and stems and produce foams. Conversely, treatments of adult trial were applied on the plants immediately before placing the insects inside the microcosm. To avoid contamination between treatments, plants were sprayed outside the climatic chamber, left to air dry for 15 min, then returned to the chamber. Treatments were applied in mid–April and mid–June 2019. A data logger placed inside the climatic chamber registered a temperature of 24.6±1.5 °C and 75.1±5.6 % RH during the nymphal experimental trial and 22.1±0.65 °C and 86.1±5.1 % RH during the adult experimental trial.

#### 2.3.3 Mortality assessment

After the treatments, microcosms were visually inspected every 2–3 days (i.e. two/three-times per week), assessing the survival rate of insects and collecting dead individuals. Newly emerging adults were also counted, and immediately removed to avoid recounting. When required to correctly assess the number of alive nymphs, foams on plant inside microcosm were gently open with a tiny brush, cleaned in 95% alcohol between each replica inspection, to avoid possible transfer of fungi between replicas. The inspections were carried out for up to three weeks or when no alive individuals were observed for two inspections in a row, allowing precise estimation of the speed of fungal strain action.

Samples of dead/symptomatic insects were collected to re-isolate the fungal strains. Dead insects were at once placed in 1.5 mL Eppendorf tubes and individually surface sterilized with a small brush dipped in a solution of 1% sodium hypochlorite and then in distilled sterile water to eliminate sodium hypochlorite residues. Individuals were subsequently stored in Petri plates fitted with moistened filter paper. Insects were incubated at 23 °C for 10 d to encourage mycelium development and daily observation was performed for the presence or absence of mycelium growth and sporulation. Once hyphal outgrowth of presumptive pathogens was detected, a first observation under the stereomicroscope provided an initial assessment of the fungal strain; then, insects were impressed on MA medium supplemented with penicillin G (0.1 mg/mL) and streptomycin (0.5 mg/mL) for fungal re-isolation. Multiplicative structures were observed under optical microscope to confirm the preliminary identification and assess if the fungi isolated from the insects were related to the inoculated species.

### 2.4 Efficacy of *L. aphanocladii* DSF on *P. spumarius* nymphs in pathogenicity bioassays

#### 2.4.1 Fungal strains and propagule preparation

Based on the results of preliminary tests, the fungal strain *L. aphanocladii* DSF was selected and used in pathogenicity bioassays against *P. spumarius* nymphs, both in conidia and blastospore formulations. In addition, *B. bassiana* ATCC 74040 strain from the commercial product Naturalis® (Biogard, Grassobbio, Bergamo, Italy) and the bio-insecticide Mycotal® (Koppert, Scarborough, Ontario, Canada) based on *Lecanicillium muscarium* Ve6 strain were used as positive controls.

*L. aphanocladii* DSF blastospores obtained from liquid substrate growth of the fungus were provided by the agrochemical company Globachem (Sint-Truiden, Belgium) and kept at 4 °C until use. They were supplied as liquid formulation or freeze-dried spores. The latter were rehydrated in sterile deionized water for 30 min at room temperature before use. Appropriate dilutions of blastospore suspensions were prepared to reach the final concentration of 1 × 10^8^ blastospore/mL on the same day of experimental treatments, and added of Tween 80 (0.01 % v/v) for a homogeneous dispersion of the propagules. *L. aphanocladii* DSF conidia suspensions were prepared at DISAFA according to the protocol described above and propagule viability was also determined as previously reported (2.3.1).

*B. bassiana* ATCC 74040 strain was isolated from Naturalis® dried powder by streak plate method to obtain a pure culture stored at 4 °C before use; afterwards, a conidia suspension was prepared and adjusted to 1 × 10^8^ conidia/mL according to the procedure described for *L. aphanocladii* DSF (2.3.1). Differently, Mycotal® was used as supplied by manufacturer (10^10^ blastospores/g) and rehydrated in sterile water immediately before use to obtain a suspension of 10^8^ blastospores/mL.

#### 2.4.2 Experimental theses and data collection

Pathogenicity bioassays on *P. spumarius* nymphs were carried out, in controlled conditions, in mid-April 2021 (trial 1, T1) and 2022 (trial 2, T2). The experimental design is reported in Table 1. Briefly, in T1 ten experimental theses were tested, six determined by the combination of fungus (*L. aphanocladii* DSF, *La*), propagules (conidia, *C* or blastospores, B) and natural adjuvants (*Ad1* and *Ad2*, provided by Globachem, composition under business secrecy) to be added to propagule suspensions in order to evaluate their biocontrol efficacy in synergy with fungal propagules; also, four control levels were tested: Mycotal (*Myc*), *adj1*, *adj2* and water (*Wtr*). In T2, *L. aphanocladii* blastospores (*LaB*) and *L. aphanocladii* conidia (*LaC*) theses were compared to *B. bassiana* Naturalis® conidia (*NCf*), used as positive control, and water (negative control, *Wtr*). All the previously described treatments were applied 24 h after placing the nymphs on plants, in order to allow nymphs to settle on leaves and stems and produce foams. Conversely, an additional thesis was tested to investigate the effect of foam on EPF effectiveness, involving nebulization of *B. bassiana* Naturalis® conidia suspension on nymphs immediately after placing them on plants, before foam formation (*NCbf*). The *V. faba* microcosms set up for the treatments are described above (2.3.2). In each microcosm, 10 nymphs of *P. spumarius* were placed, carefully took with brush and placed onto plant leaves from maintenance rearing present at DISAFA. Each thesis was replicated in three (T1) or five (T2) microcosms, for a total of 30– 50 individuals (in average 2^nd^–3^rd^ instar nymphs) for treatment. A data logger placed inside climatic chamber registered hourly climate data for each trial during the experimental periods (mean±SD – 2021: 22.8±1.1 °C and 79.2±11.6 % RH; 2022: 23.4±1.6 °C and 73.5±7.6 % RH).

**Table 1:** Experimental design of trial 1 (T1) in 2021.

| Code | Fungus | Propagule* | Adjuvant | Concentration<br>(propagule/mL) | Replicas | Nymphs/<br>replica |
| --- | --- | --- | --- | --- | --- | --- |
| **TRIAL 1 - 2021** |  |  |  |  |  |  |
| LaB_Ad1 | *L. aphanocladii* | Blastospores | Adjuvant 1 + Tween 80 | $1 \times 10^8$ | 3 | 10 |
| LaB_Ad2 | *L. aphanocladii* | Blastospores | Adjuvant 2 + Tween 80 | $1 \times 10^8$ | 3 | 10 |
| LaB_Wtr | *L. aphanocladii* | Blastospores | Tween 80 | $1 \times 10^8$ | 3 | 10 |
| LaC_Ad1 | *L. aphanocladii* | Conidia | Adjuvant 1 + Tween 80 | $1 \times 10^8$ | 3 | 10 |
| LaC_Ad2 | *L. aphanocladii* | Conidia | Adjuvant 2 + Tween 80 | $1 \times 10^8$ | 3 | 10 |
| LaC_Wtr | *L. aphanocladii* | Conidia | Tween 80 | $1 \times 10^8$ | 3 | 10 |
| Myc | *L. muscarium* Ve6<br>(Mycotal) | Conidia | — | $1 \times 10^8$ | 3 | 10 |
| Ad1 | — | — | Adjuvant 1 | $1 \times 10^8$ | 3 | 10 |
| Ad2 | — | — | Adjuvant 2 | $1 \times 10^8$ | 3 | 10 |
| Wtr<sup>+</sup> | — | — | Tween 80 | - | 3 | 10 |
| **TRIAL 2 - 2022** |  |  |  |  |  |  |
| LaB | *L. aphanocladii* | Blastospores | Tween 80 | $1 \times 10^8$ | 5 | 10 |
| LaC | *L. aphanocladii* | Conidia | Tween 80 | $1 \times 10^8$ | 5 | 10 |
| NCf | *B. bassiana* ATCC<br>74040 (Naturalis) | Conidia | Tween 80 | $1 \times 10^8$ | 5 | 10 |
| NCbf | *B. bassiana* ATCC<br>74040 (Naturalis) | Conidia | Tween 80 | $1 \times 10^8$ | 5 | 10 |
| Wtr<sup>+</sup> | — | — | Tween 80 | - | 5 | 10 |

\* blastospores were supplied by Globachem as liquid formulation in Trial 1 and freeze dried spores in Trial 2
<sup>+</sup> water treatment: negative control

After the treatments, mortality assessment was performed as previously reported (2.3.3). Dead insects were surface sterilized, as previously reported, and incubated in Petri plates containing filter paper moistened with sterile distilled water to encourage mycelium development and fungal re-isolation on culture medium was performed to assess if the fungi isolated from the cadavers were attributable to *L. aphanocladii* DSF, *B. bassiana* ATCC 74040 and *L. muscarium* Ve6 strains.

### 2.5 Statistical analysis

The estimates of nymphal survival/mortality rates were calculated using the counts of both the dead nymphs and the adults emerging, being the latter a more reliable proxy of the survival of individuals, given the difficulty in correctly counting the nymphal stages inside the microcosm. The influence of experimental treatment on survival of *P. spumarius* nymphs was hence inferred by both nymphal mortality (i.e. proportion of nymphs found dead) and adult stage emergence (i.e. total number of adults on total number of nymphs placed into each microcosm before the treatment). The two dependent variables were modelled as end point data – i.e. nymphal mortality or emergence proportion during the entire duration of trials – using binomial GLMs (glm function in package *stat*). When appropriate, mean bias-reducing adjusted scores approach was used (function brglmFit in *brglm2* package, Firth 1993) to avoid quasi–complete separation due to a category (Thesis) with all (1) or no (0) nymphs dead or emerged. The probability of nymphal status (alive, dead, emerged as adult) through time post treatment as an effect of experimental treatment was analysed by multi-state Cox proportional hazards models (coxph function in package *survival*, Therneau et al. 2022), with nymphs not found during visual inspection – possibly escaped – as censoring events. Data were tested to confirm they fit model assumptions using the coxzph function (*survival* package). The end-point proportion of adult stage survival was modelled using a binomial GLM (glm function in package *stats*). All analyses were performed and plots produced using the software R (R Core Team 2020).

## 3. RESULTS

### 3.1 Identification of EPF strains isolated from *P. spumarius*

Four fungal isolates were identified based on rDNA-ITS homology analysis. ITS sequences showed 100% similarity to *Lecanicillium aphanocladii* (GenBank sequence ID: KC574075.1), 99,8% similarity to *Beauveria bassiana* (GenBank sequence ID: MF872381.1), 99,6 % similarity to *Fusarium equiseti* (GenBank sequence ID: MN559437.1) and 98% similarity to *Conidiobolus coronatus* (GenBank sequence ID: AJ345094.1). Molecular identification was supported by macroscopic and microscopic analysis of phenotypic characteristics as colony colour, hyphal aspect, multiplicative structure and propagule morphology.

For all the species identified, literature have reported evidence on their entomopathogenic potential (Souza et al. 2014; Sharma and Marques 2018; Wronska et al. 2018; Zhao et al. 2023). Thus, the four fungal strains were studied in preliminary trials to test their efficacy against the target insect *P. spumarius*.

### 3.2 Pathogenic activity of EPF isolates against *P. spumarius* nymphs and adults

#### Nymphs

The viability of conidia used in the tests was comparable between the different strains: 97.2 ± 2.2% for *B. bassiana*, 95.8 ± 4.0% for *L. aphanocladii*, 93.1 ± 2.5% for *C. coronatus* and 90.2 ± 4.3% for *F. equiseti*.

The efficacy of the four EPF strains on *P. spumarius* nymphs (i.e. end-point mortality percentage) was significantly different. Indeed, the probability of nymphal death was significantly higher for individuals exposed to *B. bassiana* (46.7%, 14/30), *L. aphanocladii* and *C. coronatus* (43.3%, n = 13/30), compared to *F. equiseti* (3.3%, 1/30) and control (*Wtr*) (0%, 0/32) (GLM: χ^2^ = 38.73, df = 4, p < 0.001) (Figure 1a; Table 2). The hazard ratio of nymphal death (i.e. average risk of death) was significantly higher for nymphs treated with *B. bassiana* (HR = 19.5, CI = 2.8–137), *C. coronatus* (HR = 20.1, CI = 2.76–146) and *L. aphanocladii* (HR = 15.9, CI = 2.13–119) than control (*Wtr*). *Fusarium equisetii* treatment did not significantly increase nymphal death risk compared to control (HR = 1.11, CI = 0.07–17.3), while it showed significantly lower death risk than the other tested EPF species (HR = 14.3–18.0) (Figure 2).

**Figure 1:**
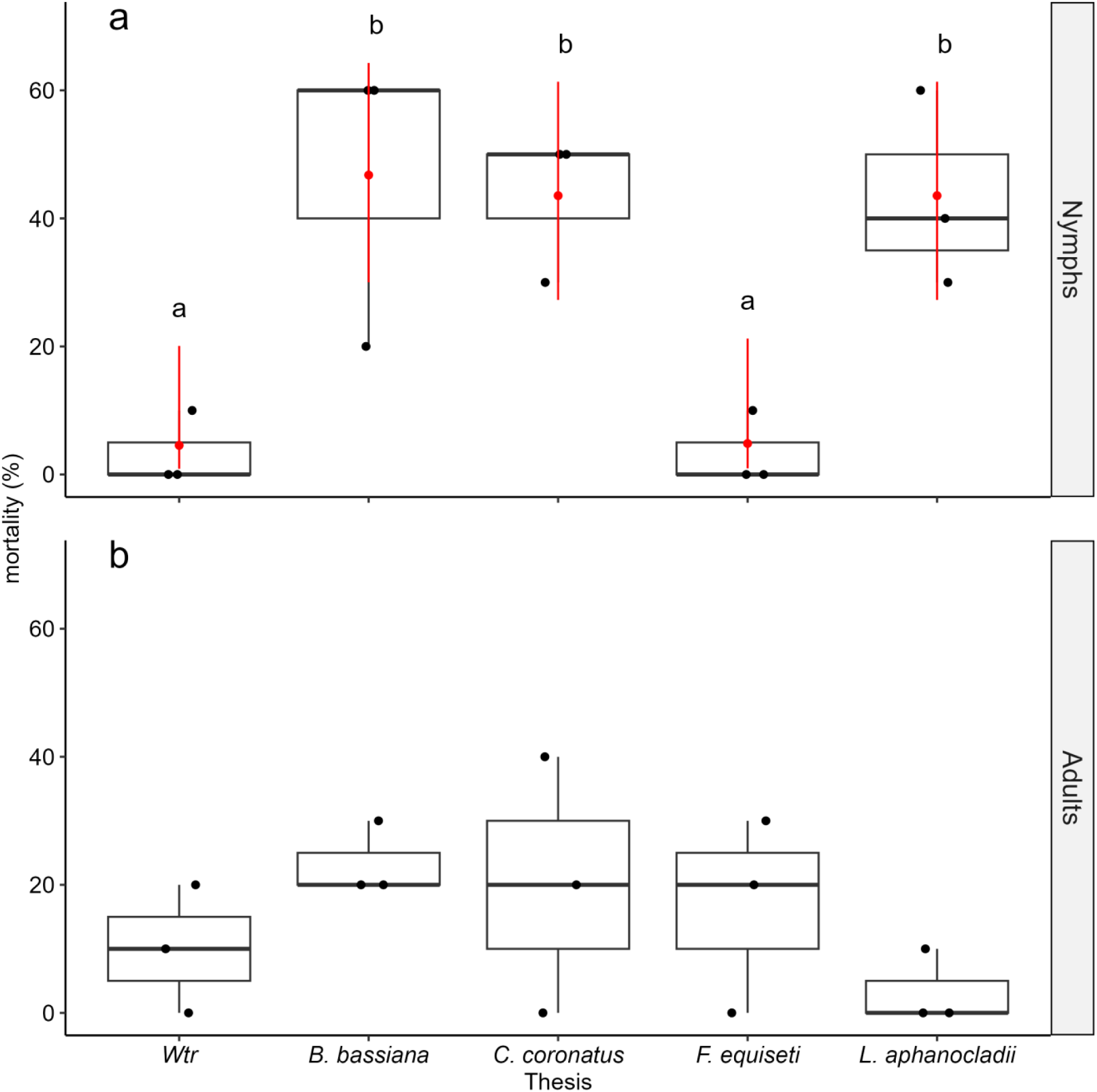
T1-2019; Percentage of mortality observed (boxplots) and predicted by binomial GLM (red point range) for nymphal and adult stages of Philaenus spumarius in experimental trials of pathogenic activity of EPF isolates. Treatments sharing the same letter within a facet are not significantly different (α = 0.05, Tukey-adjusted). Predictions by binomial GLM are not represented for adult stage because the effect of thesis on mortality rate was not significant, i.e. no differences in mortality rate among experimental theses.

**Figure 2:**
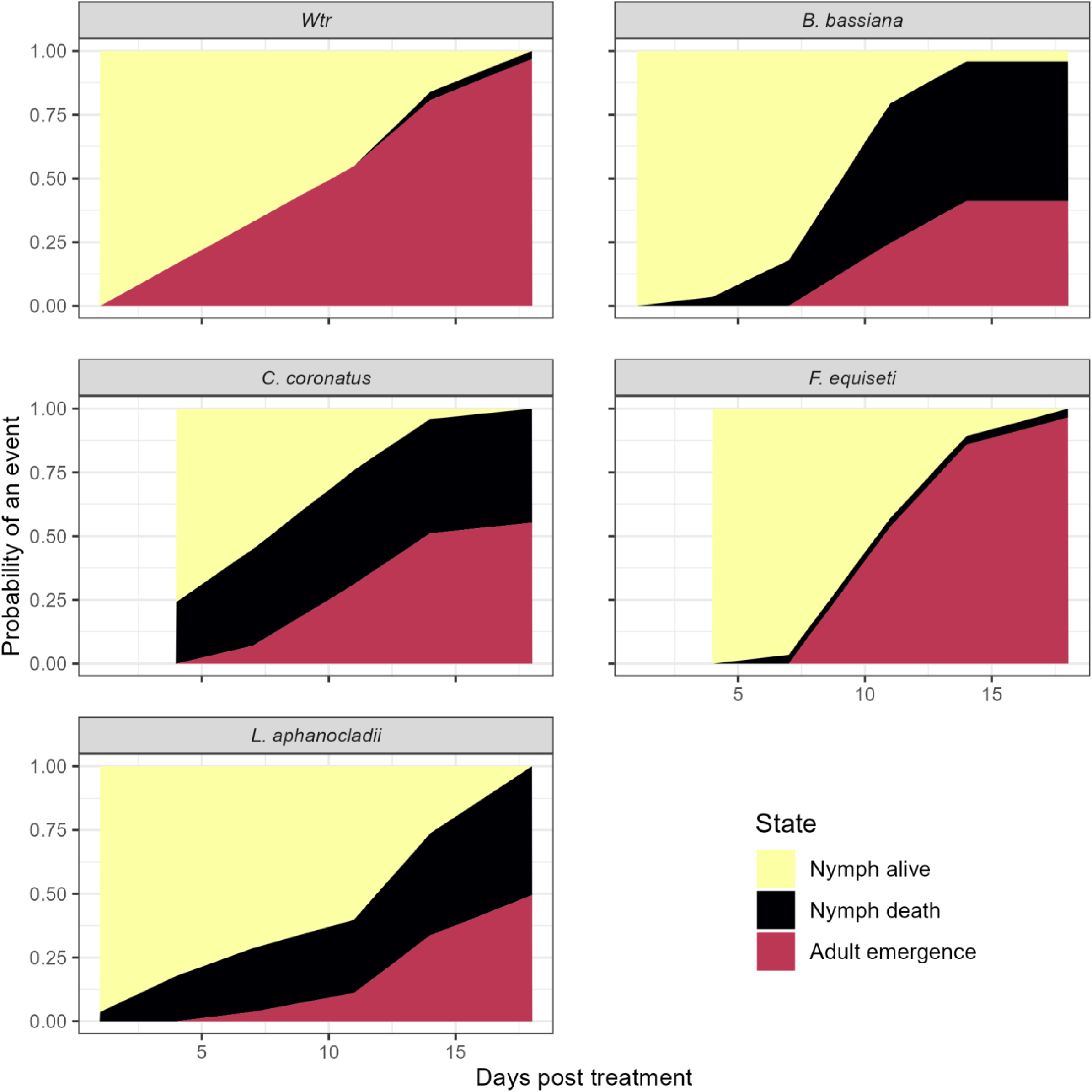
T1-2019; Survival curves fitted by multi-state Cox Hazard Model. Areas of different colors represent the estimated probability of each P. spumarius nymphal state (alive, death, adult emergence) over time after the experimental treatment.

**Table 2:** Pathogenic activity of naturally occurring EPF strains against Philaenus spumarius nymphs. Nymphal mortality and adult emergence estimates (%) and variance [standard error (SE), lower and upper confidence limit 95% (UCL)] calculated by binomial GLM (actual results reported in the text). Theses sharing the same letter in group column are not significantly different (α = 0.05, Tukey-adjusted).

| Variable | Thesis | Estimate (%) | SE | LCL | UCL | group |
| --- | --- | --- | --- | --- | --- | --- |
| **Nymphal mortality** |  |  |  |  |  |  |
|  | *Wtr* | 4.55 | 3.68 | 0.89 | 20.08 | a |
|  | *F. equiseti* | 4.84 | 3.92 | 0.95 | 21.23 | a |
|  | *C. coronatus* | 43.55 | 9.05 | 27.26 | 61.35 | b |
|  | *L. aphanocladii* | 43.55 | 9.05 | 27.26 | 61.35 | b |
|  | *B. bassiana* | 46.77 | 9.11 | 30.02 | 64.29 | b |
| **Adult emergence** |  |  |  |  |  |  |
|  | *Wtr* | 87.50 | 71.06 | 95.23 | b | b |
|  | *F. equiseti* | 86.67 | 69.41 | 94.9 | b | b |
|  | *C. coronatus* | 50 | 32.83 | 67.17 | a | a |
|  | *L. aphanocladii* | 40 | 24.31 | 58.05 | a | a |
|  | *B. bassiana* | 33.33 | 18.97 | 51.65 | a | a |

The probability of adult emergence was significantly lower for individuals exposed to *B. bassiana* (33.3%, 10/30), *L. aphanocladii* (40%, n = 12/30) and *C. coronatus* (50%, n = 15/30), compared to *F. equiseti* (86.7%, 26/30) and control (water) (87.5%, 28/32) (GLM: χ^2^ = 36.92, df = 4, p < 0.001) (Table 2). Thus, the hazard ratio of adult emergence was significantly lower for individuals treated with *B. bassiana* (HR = 0.69, CI = 0.5–0.96) and *L. aphanocladii* (HR = 0.47, CI = 0.40–0.55) compared to control (*Wtr*). No differences in adult emergence risk were observed between *C. coronatus* (HR = 1.14, CI = 0.94–1.37) and *F. equiseti* (HR = 1.1, CI = 0.82–1.46) compared to the control thesis (*Wtr*) (Figure 2).

#### Adults

No significant differences in mortality of adult stage of *P. spumarius* were observed among the four tested EPF strains and compared to the control (*Wtr*) (χ^2^ = 7.14, df = 4, p = 0.13) (Figure 1b).

For the subsequent pathogenicity bioassays focused on *P. spumarius* nymphs, *L. aphanocladii* DSF strain was selected among the others because judged the most promising in terms of performance and noteworthy for the novelty of the species it belongs to. The strain was named *L. aphanocladii* DSF due to its isolation at Department of Agricultural, Forest and Food Sciences (DISAFA) laboratories.

Despite the performance comparable to *L. aphanocladii*, *C. coronatus* and *B. bassiana* strains were not preferred because the latter belongs to a species already well known and present on the market for biocontrol, and the former can cause mycosis in mammals, including humans. Actually, its incidence as a human pathogen remains low but it may represent a risk for immunocompromised individuals (Vilela and Mendoza 2018).

### 3.3 L. aphanocladii DSF efficacy on P. spumarius nymphs

The pathogenicity of *L. aphanocladii* DSF against *P. spumarius* nymphs was assessed under controlled conditions in two separate trials (T1 and T2). The viability of fungal propagules was checked immediately before their use and was of 95.5 ± 4.2% and 97.1 ± 2.2% for *L. aphanocladii* blastospores used in T1 and T2, respectively; 93.8 ± 3.3% and 94.8 ± 4.3% for *L. aphanocladii* conidia in T1 and T2, respectively; 97.4 ± 2.0 % for *B. bassiana* Naturalis® conidia used in T2.

#### T1

The end-point nymphal mortality (i.e., probability of death of a nymph) was overall significantly different among experimental theses (GLM: χ^2^ = 44.08, df = 9, p < 0.001), with the EPF-treated microcosms, except for *LaC_Ad1* thesis, showing higher mortality rates compared to those treated with water (control). The highest nymphal mortality rates were recorded in *LaB_Ad2* thesis (63.3 ± 8.8%), although not significantly higher than the other EPF theses (ranging 44.8–60.0%), again except for *LaC_Ad1* one (18.5 ± 7.5%) (Figure 3 and Table 3). Mycotal *Myc* (16.7 ± 6.8%) and the adjuvant 1 *Ad1* (6.9 ± 3.3%) caused significantly less mortality in

**Figure 3:**
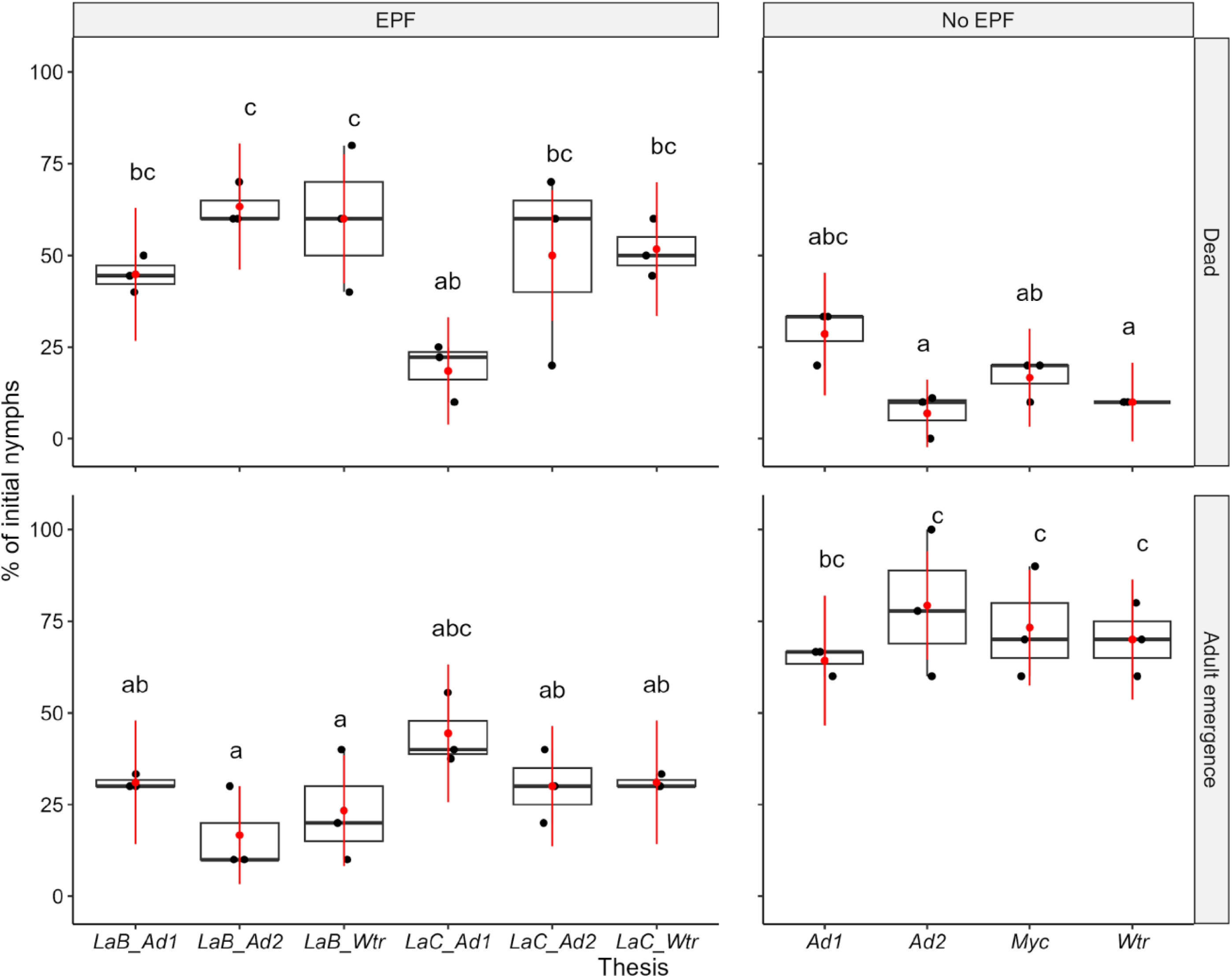
T1-2021; Percentage of initial nymphs of Philaenus spumarius found dead or emerged as adults during the 2021 experimental trial (T1) of Lecanicillium aphanocladii DSF efficacy. Data observed (boxplots) and predicted by binomial GLM (red point range) are reported, and treatments sharing the same letter within a row are not significantly different (α = 0.05, Tukey-adjusted).

**Table 3:** T1-2021; Lecanicillium aphanocladii DSF efficacy on Philaenus spumarius nymphs in 2021 trial (T1). Nymphal mortality and adult emergence estimates (%) and variance [standard error (SE), lower and upper confidence limit 95% (UCL)] calculated by binomial GLM. Theses sharing the same letter in group column are not significantly different (α = 0.05, Tukey-adjusted).

| Variable | Thesis | Estimate (%) | SE | LCL | UCL | group |
| --- | --- | --- | --- | --- | --- | --- |
| **Nymphal mortality** |  |  |  |  |  |  |
|  | Ad2 | 6.90 | 4.71 | 0 | 16.12 | a |
|  | Wtr | 10.00 | 5.48 | 0 | 20.74 | a |
|  | Myc | 16.67 | 6.80 | 3.33 | 30.00 | ab |
|  | LaC_Ad1 | 18.52 | 7.48 | 3.87 | 33.17 | ab |
|  | Ad1 | 28.57 | 8.54 | 11.84 | 45.30 | abc |
|  | LaB_Ad1 | 44.83 | 9.23 | 26.73 | 62.93 | bc |
|  | LaC_Ad2 | 50.00 | 9.13 | 32.11 | 67.89 | bc |
|  | LaC_Wtr | 51.72 | 9.28 | 33.54 | 69.91 | bc |
|  | LaB_Wtr | 60.00 | 8.94 | 42.47 | 77.53 | c |
|  | LaB_Ad2 | 63.33 | 8.80 | 46.09 | 80.58 | c |
| **Adult emergence** |  |  |  |  |  |  |
|  | LaB_Ad2 | 16.67 | 6.80 | 3.33 | 30.00 | a |
|  | LaB_Wtr | 23.33 | 7.72 | 8.20 | 38.47 | a |
|  | LaC_Ad2 | 30.00 | 8.37 | 13.60 | 46.40 | ab |
|  | LaB_Ad1 | 31.03 | 8.59 | 14.20 | 47.87 | ab |
|  | LaC_Wtr | 31.03 | 8.59 | 14.20 | 47.87 | ab |
|  | LaC_Ad1 | 44.44 | 9.56 | 25.70 | 63.19 | abc |
|  | Ad1 | 64.29 | 9.06 | 46.54 | 82.03 | bc |
|  | Wtr | 70.00 | 8.37 | 53.60 | 86.40 | c |
|  | Myc | 73.33 | 8.07 | 57.51 | 89.16 | c |
|  | Ad2 | 79.31 | 7.52 | 64.57 | 94.05 | c |

*P. spumarius* nymphs than *L. aphanocladii* formulations, comparable to that observed in control thesis *Wtr* (10.0 ± 0.0%). The treatment applied was indeed a significant predictor of nymphal risk of death. Nymphs treated with *L. aphanocladii* formulations were significantly more likely to die compared to control (*Wtr*) (HR EPF treated:control = 4.79–7.94) (Figure 4; Table 4). The only exception was *LaC_Ad1* thesis, that did not cause a significant increase in the probability of nymphal death (HR = 2.39, CI = 0.56–10.19). Mycotal did not significantly increase the nymphal death likelihood compared to control (*Wtr*) (HR = 1.6, CI = 0.35–7.31) (Figure 4; Table 4).

**Figure 4:**
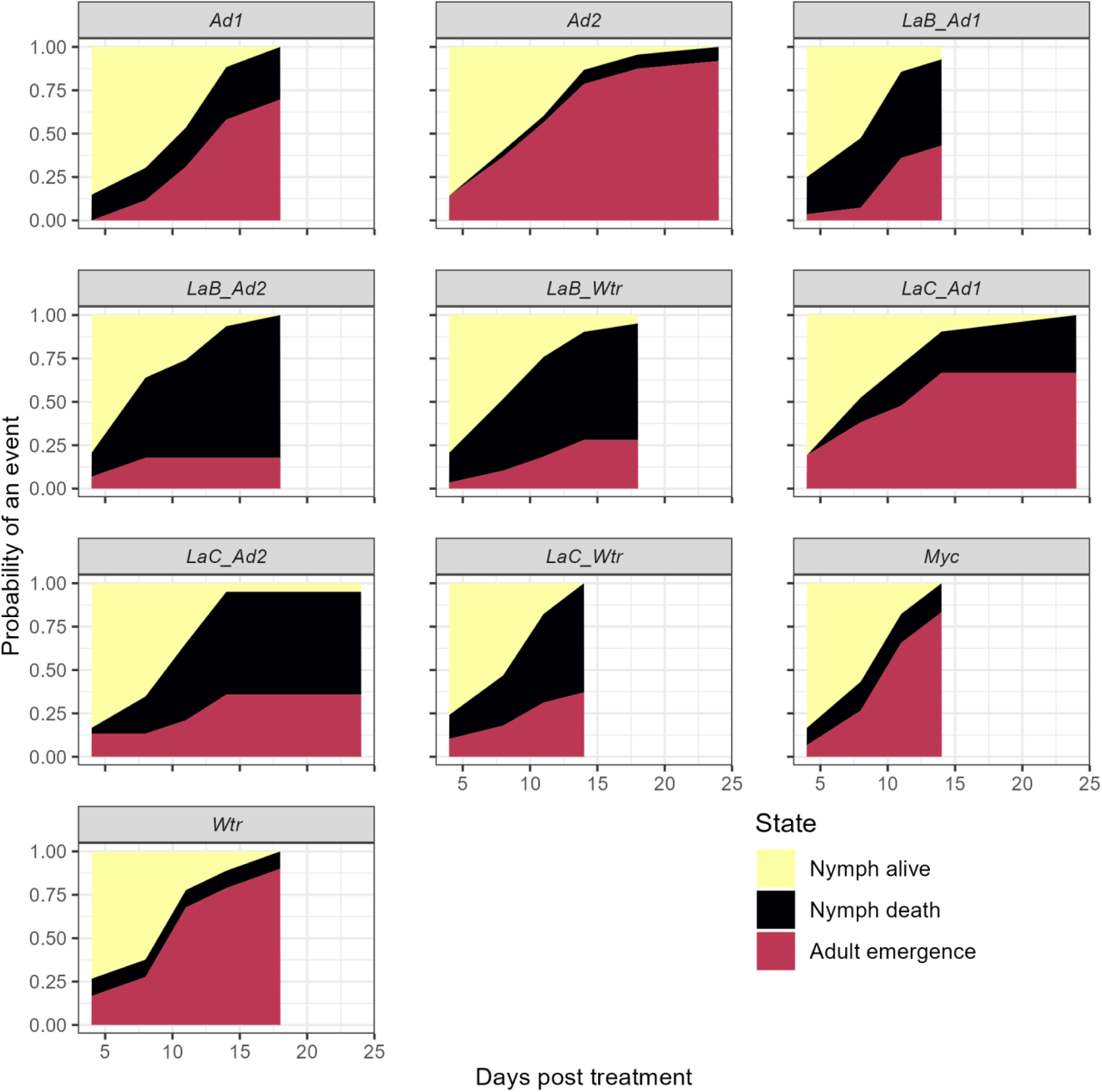
T1-2021; Survival curves of nymphs Philaenus spumarius treated with different Lecanicillium aphanocladii formulations in trial T1, fitted by multi-state Cox Hazard Model. Areas of different colors represent the estimated probability of each P. spumarius nymphal state (alive, death, adult emergence) over time after the experimental treatment.

**Table 4:** T1-2021; Hazard ratios and 95% confidence intervals (95% CI) for transitioning of Philaenus spumarius nymphs from alive to dead and alive to adult emergence for experimental theses respect to control (Wtr) and LaB_Ad2, representing minimum and maximum mortality, respectively. Comparisons among hazard ratios of transition states for all the experimental theses can be found in Supplemetary Materials.

| Transition | Treatment | With respect to | Hazard ratio | 95% CI |  | z | p-value |  |
| --- | --- | --- | --- | --- | --- | --- | --- | --- |
| Alive to dead | Ad1 | Wtr | 2.06 | 0.51 | 8.39 | 1.01 | 0.31 |  |
|  | Ad2 | Wtr | 0.59 | 0.10 | 3.58 | -0.58 | 0.56 |  |
|  | LaB_Ad1 | Wtr | 5.77 | 1.51 | 22.13 | 2.56 | 0.01 | ** |
|  | LaB_Ad2 | Wtr | 7.94 | 2.21 | 28.59 | 3.17 | 0.00 | *** |
|  | LaB_Wtr | Wtr | 6.30 | 1.72 | 23.04 | 2.78 | 0.01 | ** |
|  | LaC_Ad1 | Wtr | 2.39 | 0.56 | 10.19 | 1.17 | 0.24 |  |
|  | LaC_Ad2 | Wtr | 4.79 | 1.34 | 17.20 | 2.40 | 0.02 | * |
|  | LaC_Wtr | Wtr | 6.05 | 1.67 | 21.90 | 2.74 | 0.01 | ** |
|  | Myc | Wtr | 1.60 | 0.35 | 7.31 | 0.60 | 0.55 |  |
|  | Wtr | LaB_Ad2 | 0.13 | 0.03 | 0.45 | -3.17 | 0.00 | *** |
|  | Ad1 | LaB_Ad2 | 0.26 | 0.11 | 0.60 | -3.15 | 0.00 | *** |
|  | Ad2 | LaB_Ad2 | 0.07 | 0.02 | 0.31 | -3.57 | 0.00 | *** |
|  | LaB_Ad1 | LaB_Ad2 | 0.73 | 0.36 | 1.48 | -0.88 | 0.38 |  |
|  | LaB_Wtr | LaB_Ad2 | 0.79 | 0.42 | 1.48 | -0.73 | 0.47 |  |
|  | LaC_Ad1 | LaB_Ad2 | 0.30 | 0.12 | 0.74 | -2.62 | 0.01 | ** |
|  | LaC_Ad2 | LaB_Ad2 | 0.60 | 0.33 | 1.09 | -1.68 | 0.09 |  |
|  | LaC_Wtr | LaB_Ad2 | 0.76 | 0.42 | 1.37 | -0.90 | 0.37 |  |
|  | Myc | LaB_Ad2 | 0.20 | 0.07 | 0.56 | -3.08 | 0.00 | *** |
| Alive to adult | Ad1 | Wtr | 0.54 | 0.32 | 0.92 | -2.29 | 0.02 | * |
|  | Ad2 | Wtr | 0.74 | 0.41 | 1.36 | -0.96 | 0.33 |  |
|  | LaB_Ad1 | Wtr | 0.52 | 0.25 | 1.08 | -1.75 | 0.08 |  |
|  | LaB_Ad2 | Wtr | 0.25 | 0.08 | 0.74 | -2.49 | 0.01 | * |
|  | LaB_Wtr | Wtr | 0.28 | 0.12 | 0.65 | -2.93 | 0.00 | ** |
|  | LaC_Ad1 | Wtr | 0.85 | 0.40 | 1.84 | -0.40 | 0.69 |  |
|  | LaC_Ad2 | Wtr | 0.30 | 0.14 | 0.66 | -3.00 | 0.00 | ** |
|  | LaC_Wtr | Wtr | 0.46 | 0.21 | 1.01 | -1.94 | 0.05 |  |
|  | Myc | Wtr | 1.11 | 0.66 | 1.86 | 0.39 | 0.70 |  |
|  | Wtr | LaB_Ad2 | 4.05 | 1.35 | 12.17 | 2.49 | 0.01 | ** |
|  | Ad1 | LaB_Ad2 | 2.19 | 0.76 | 6.31 | 1.45 | 0.15 |  |
|  | Ad2 | LaB_Ad2 | 3.01 | 0.99 | 9.21 | 1.93 | 0.05 | * |
|  | LaB_Ad1 | LaB_Ad2 | 2.12 | 0.65 | 6.86 | 1.25 | 0.21 |  |
|  | LaB_Wtr | LaB_Ad2 | 1.12 | 0.32 | 3.96 | 0.18 | 0.86 |  |
|  | LaC_Ad1 | LaB_Ad2 | 3.46 | 1.01 | 11.85 | 1.98 | 0.05 | * |
|  | LaC_Ad2 | LaB_Ad2 | 1.23 | 0.36 | 4.16 | 0.34 | 0.74 |  |
|  | LaC_Wtr | LaB_Ad2 | 1.85 | 0.55 | 6.25 | 0.99 | 0.32 |  |
|  | Myc | LaB_Ad2 | 4.48 | 1.56 | 12.91 | 2.78 | 0.01 | ** |

The adult emergence rate was also significantly different among the tested theses, with an expected inverse trend compared to nymphal mortality (GLM: χ^2^ = 51.29, df = 9, p < 0.001). All the EPF-treated theses showed significantly lower emergence rates (16.7–31.0%) than *Wtr* (70.0 ± 0.0%), *Myc* (73.3 ± 8.1%), and *Ad1* (79.3 ± 7.5%) ones (Figure 3 and Table 3). The results obtained from the end-point adult emergence were confirmed by the over time competing risks analysis on the status of nymphs. Nymphs treated with *LaB_Ad2*, *LaB_Wtr*, *LaC_Ad2* were significantly less likely to become adult than those in control (HR EPF treated:control = 0.25 – 0.54) (Figure 4; Table 4). Also *Ad1* thesis decreased the likelihood of adult emergence (HR = 0.54, CI = 0.32– 0.92), while neither *LaC_Ad1* nor *Myc* treatments significantly reduced the adult emergence rate (HR *LaC_Ad1*:control = 0.85, CI = 0.4–1.84; HR *Myc*:control = 1.11, CI = 0.66–1.86).

A total of 156 dead nymphs were collected during the visual inspections, and all were prepared for fungal reisolation. *L. aphanocladii* was successfully reisolated from approximately half of them (n = 65, 47.8 %), all collected from fungal theses. A single dead nymph collected from Mycotal-treated microcosm did not develop *L. muscarium* mycelium.

#### T2

The end-point nymphal mortality was significantly different among experimental theses (GLM: χ^2^ = 41.77, df = 4, p < 0.001), with all the EPF-treated microcosms showing higher mortality rates of nymphs compared to those treated with water (control) (Table 4). Namely, nymphs sprayed with *L. aphanocladii* blastospores (*LaB*) showed the highest mortality rate (91.1 ± 4.3%, 47 dead/52 exposed) and were significantly more likely to die during the assay than both control (*Wtr*) (16.6 ± 6.0%, 9/51) and *L. aphanocladii* conidia (LaC) (59.3 ± 8.4%, 32/54) (Table 5). The two *B. bassiana* theses (*NCf*: 81.4 ± 6.4%, 44/55; *NCbf*: 81.3 ± 6.4%, 41/51) did not significantly differ for the nymphal mortality rate among them and from both *L. aphanocladii* formulations (*LaB* and *LaC*) (Figure 5). The hazard ratios of death were significantly higher for all the individuals treated with EPF strains compared to control (*Wtr*) (HR EPF-treated:control = 4.9–14.1) (Table 4), although *LaC*-treated nymphs were significantly less likely to die than those treated with the other fungal propagules (HR LaC:other EPF theses = 0.30-0.44) (Table 3). The progression of nymphal mortality was significantly different among experimental theses, with nymphs treated with EPFs reaching 50% nymphal mortality within 3–4 days post treatment (CI 95% 1–4 days), except for *LaC* that reached 50 % mortality significantly later (7 days post treatment, CI 95% 7–9 days) (Figure 6).

**Figure 5:**
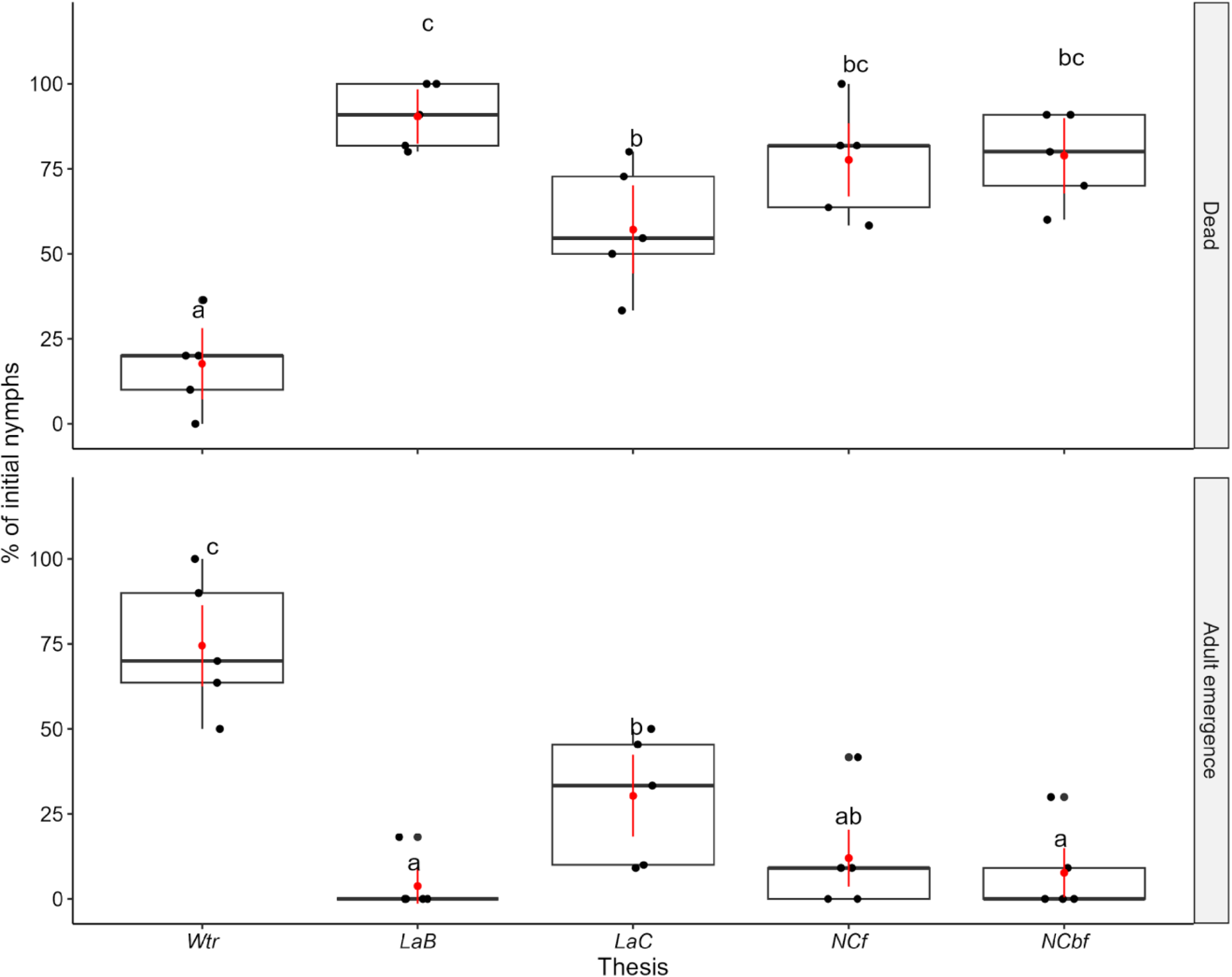
T2-2022; Percentage of initial nymphs of Philaenus spumarius found dead or emerged as adults during the 2022 experimental trial (T2) of Lecanicillium aphanocladii DSF efficacy. Data observed (boxplots) and predicted by binomial GLM (red point range) are reported, and theses sharing the same letter within a row are not significantly different (α = 0.05, Tukey-adjusted).

**Figure 6:**
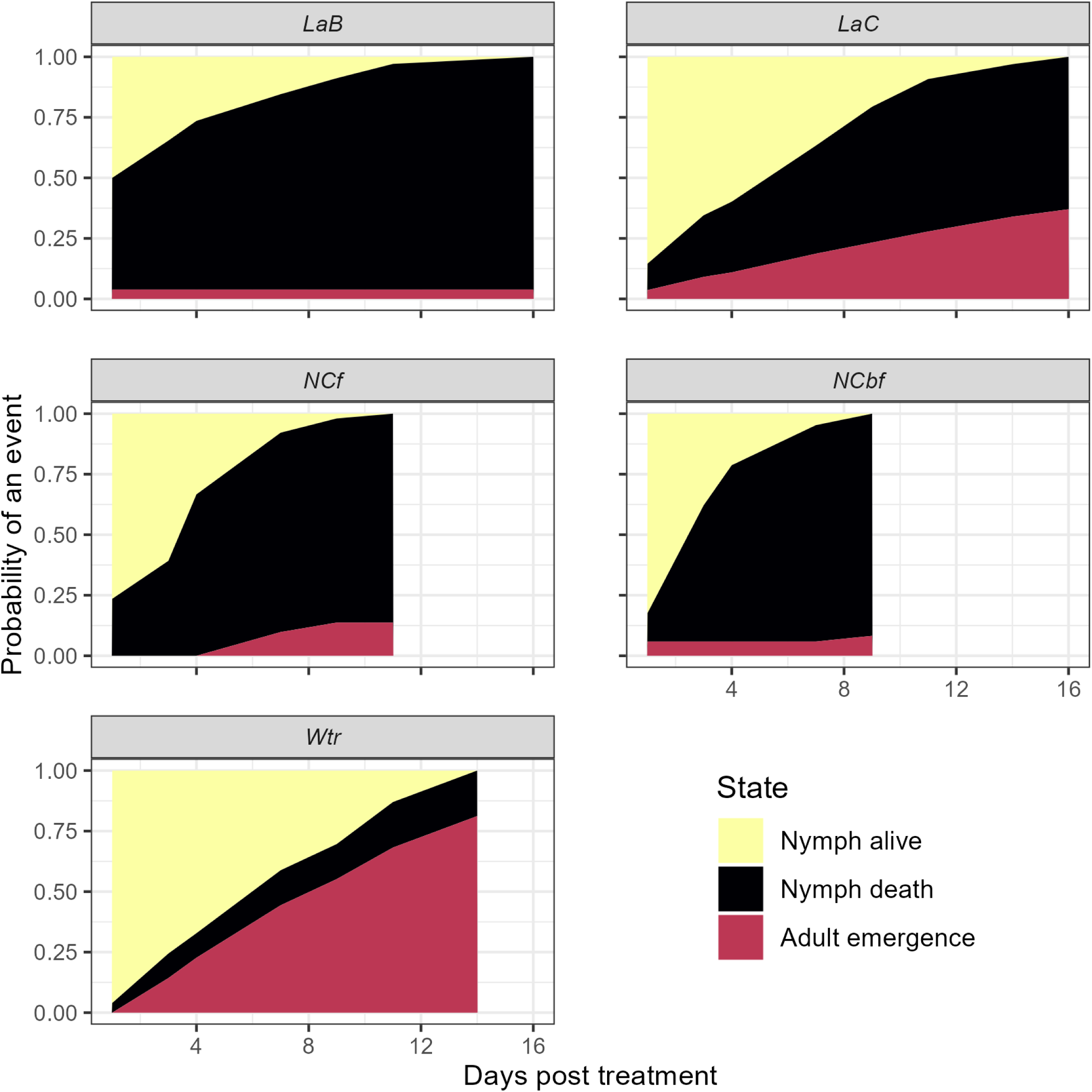
T2-2022; Survival curves of nymphs Philaenus spumarius treated with different Lecanicillium aphanocladii formulations in trial T2, fitted by multi-state Cox Hazard Model. Areas of different colors represent the estimated probability of each P. spumarius nymphal state (alive, death, adult emergence) over time after the experimental treatment.

**Table 5:** T2-2022; Lecanicillium aphanocladii DSF efficacy on Philaenus spumarius nymphs in 2022 trial (T2). Nymphal mortality and adult emergence estimates (%) and variance [standard error (SE), lower and upper confidence limit 95% (UCL)] calculated by binomial GLM. Theses sharing the same letter in group column are not significantly different (α = 0.05, Tukey-adjusted).

| Variable | Thesis | Estimate (%) | SE | LCL | UCL | group |
| --- | --- | --- | --- | --- | --- | --- |
| **Nymphal mortality** |  |  |  |  |  |  |
|  | *Wtr* | 17.65 | 5.34 | 7.18 | 28.11 | a |
|  | *LaC* | 57.14 | 6.61 | 44.18 | 70.10 | b |
|  | *NCf* | 77.59 | 5.48 | 66.85 | 88.32 | bc |
|  | *NCbf* | 78.85 | 5.66 | 67.75 | 89.95 | bc |
|  | *LaB* | 90.38 | 4.09 | 82.37 | 98.40 | c |
| **Adult emergence** |  |  |  |  |  |  |
|  | *Wtr* | 74.51 | 6.10 | 62.55 | 86.47 | c |
|  | *LaC* | 30.36 | 6.14 | 18.31 | 42.40 | b |
|  | *NCf* | 12.07 | 4.28 | 3.69 | 20.45 | ab |
|  | *NCbf* | 7.69 | 3.70 | 0.45 | 14.93 | a |
|  | *LaB* | 3.85 | 2.67 | -1.38 | 9.07 | a |

As expected, the probability of adult emergence followed an inverse pattern, hence significantly lower for individuals exposed to all EPF theses [*LaB* (3.8±4.1%, 2/47), *NCbf* (7.8±5.8%, n = 4/51), *NCf* (12.7±6.9%, 7/55) and *LaC* (31.5±9.7%, n = 17/54)] compared to control (*Wtr*) (74.5±9.4%, 38/51) (GLM: χ^2^ = 37.63, df = 4, p < 0.001) (Figure 5 and Table 5). *LaB* and *NCbf* adult emergence rates were also significantly lower than *LaC*. The over time competing risks analysis on the status of nymphs confirmed that nymphs were less likely to emerge as adults when treated with EPF thesis, in particular *LaB* one (HR *LaB:control* = 0.1, CI = 0.02–0.47) (Table 4). The progression of adult emergence through time post treatment highlighted that the emergence rate was quite constant, although different, for both *Wtr* and *LaC* theses, while for *LaB*, *NCf* and *NCbf* the few adults were emerging immediately after and within one week from the treatment (Figure 6).

One-hundred-eighty dead nymphs were collected during the visual inspections, 77.2% of which developed mycelium referable to the inoculated fungal species. Ten dead nymphs were collected from negative control (*Wtr*), and did not show any fungal growth. The symptoms of nymphs affected by mycosis due to *L. aphanocladii* DSF or *B. bassiana* Naturalis® were clearly visible under the stereomicroscope (Figure 7). *B. bassiana* Naturalis® emerging from the cadaver covered the insect with a white and rather compact mycelium while *L. aphanocladii* DSF formed a loose, cobweb-like mycelium of an off-white colour.

**Figure 7:**
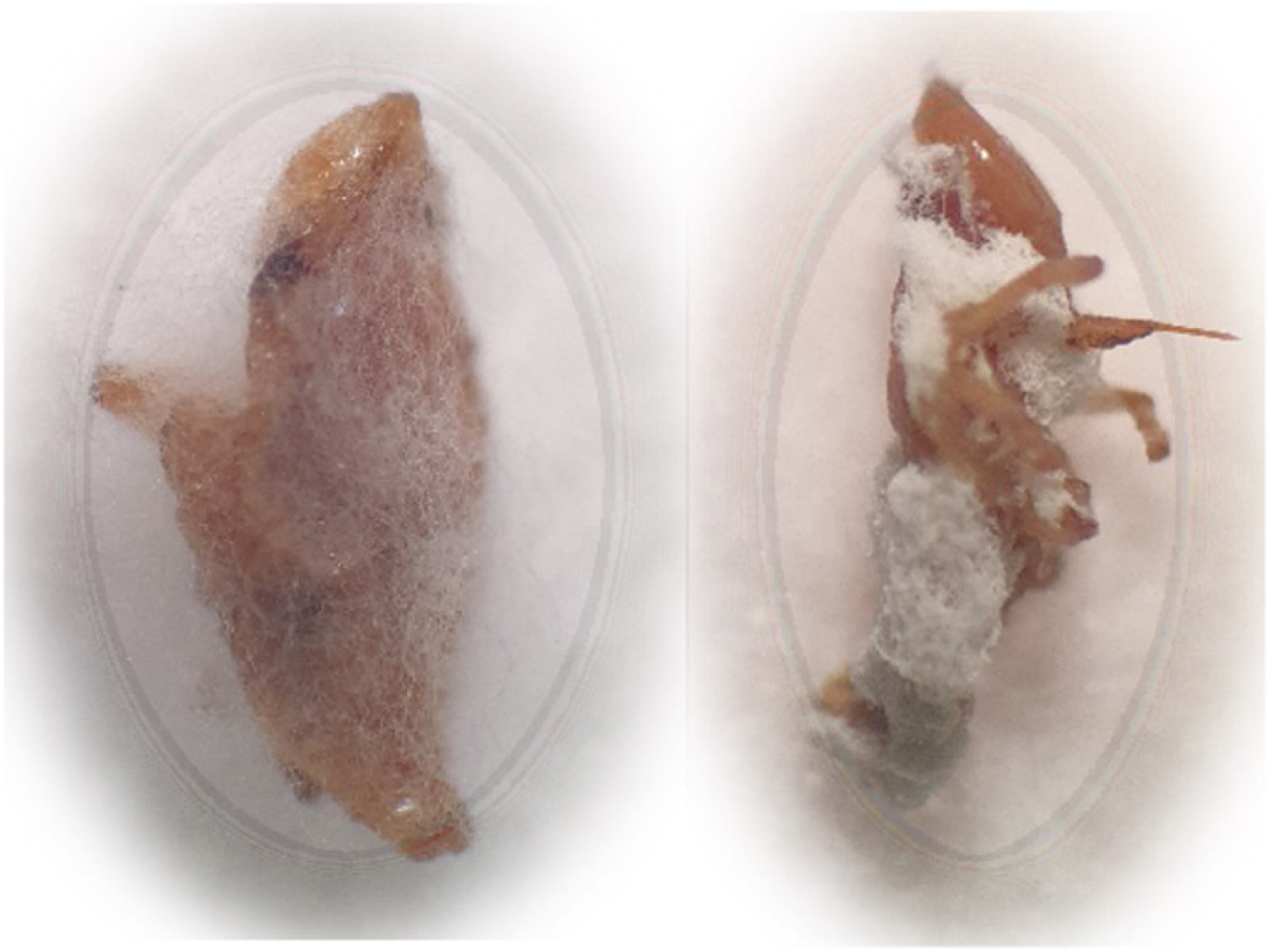
Lecanicillium aphanocladii (a) and Beauveria bassiana (b) mycelium emerging from dead Philaenus spumarius nymphs collected in trial T2.

## 4. DISCUSSION

Entomopathogenic fungi (EPF) play an important role in the natural regulation of insect populations and they have been recognized as important agents in biocontrol strategy.

The present research represents one of the first work on EPF genera and species naturally attacking *P. spumarius*, vector of the noxious phytopathogenic bacterium *X. fastidiosa*. We successfully isolated four different strains of EPF naturally infecting *P. spumarius* in the Piemonte Region (Northwestern Italy), and belonging to the species *L. aphanocladii*, *B. bassiana*, *C. coronatus* and *F. equiseti*. Preliminary experimental trials involving direct nebulization of conidia on nymphal stages, on a single potted plant, showed promising pathogenic activity of the isolates (43–46% mortality), except for *F. equiseti* (3% mortality). Conversely, preliminary trials on adults of *P. spumarius* revealed very low mortality rates of EPF-treated individuals and no significant differences compared to those exposed to control treatment (water). Similar differences in EPF lethal activity against nymphal and adult stage were not observed in previous studies involving sharpshooters (Cicadellidae: Proconiini) (Lietze et al. 2011; Cabanillas and Jones 2013), although differences in setup, target species, and fungal strains could play a role in the different outcomes. Notably, the observed different pathogenicity level of fungal strains against *P. spumarius* nymphs and adults could be influenced also by the different spraying method of propagules employed (nymphs: direct, adult: indirect), due to logistic difficulties given by the different mobility of the insect stages. This may have reduced the probability of propagule-insect contact.

Given the low efficiency against adult stage of the isolated fungal strains, subsequent bioassays were performed on nymphal stages, as a more suitable insect-stage target. Furthermore, the choice to focus on preimaginal stages was done also considering the epidemiological roles of the different life stages of *P. spumarius*. Nymphs have no relevant role in *Xf* transmission to the crop, and all the mortality caused to this stage will likely lower the adult densities in olive groves and thus the *Xf* spread likelihood (Saponari et al. 2019). Furthermore, given that *P. spumarius* adults may acquire and inoculate Xf soon after colonization of olive trees (Cornara et al. 2017; Bodino et al. 2021), even timely EPF treatments on adult stage could be not very effective in reducing *Xf* spread on the crop. Also, nymphs are characterised by low mobility, high densities in a single vegetation compartment (herbaceous cover), and they are present during a relatively humid and cool period of the year (early Spring) (Morente et al. 2018; Bodino et al. 2019). These traits make nymphs likely the best biological stage for application of EPF.

Among fungal isolates*, L. aphanocladii* DSF was considered the most promising strain to be further investigated because i) showed low rates of nymphal survival (43%) and adult emergence (40%), similar to those observed in spittlebugs treated with *B. bassiana*, a fungal species well recognized as an essential source of myco-pesticides used for numerous insect pest all over the word (Soliman et al., 2022), ii) presented an intense sporulation activity of both conidia and blastospores and iii) is a new species for the biological product market, although already found in nature as an entomopathogenic species (Zhou et al., 2020; Nedveckyte et al., 2021).

*Lecanicillium* sp., along with *Beauveria* sp., *Metarhizium* sp. and *Isaria* sp., are natural enemies of insect pests and widely studied for use in biocontrol and reducing the need for synthetic pesticides. Representative strains of the genus *Lecanicillium* are already used as active ingredients of several mycoinsecticides (Goettel et al., 2008; Bamisile et al., 2021). Totally, about 15 *Lecanicillium* spp.-based commercial preparations have been developed (Faria and Wraight, 2007), as Mycotal® and Lecatech®, formulated with *L. muscarium* and *Lecanicillium lecanii*, respectively. Currently, no products based on *L. aphanocladii* propagules are present on the market.

In nature, this species was initially isolated from fungi, such as *Agaricus* sp. and *Sphaerotheca* sp. (Zare and Gams, 2001), and insects, as mosquito larvae, *Bombyx mori* (L.) and *Trialeurodes vaporariorum* Westwood (Zare and Gams, 2001); its geographical distribution is considered cosmopolitan. More recently, *L. aphanocladii* has been found affecting the adult stage of the leaf-miner horse-chestnut pest *Cameraria ohridella* Descha & Dimic and it showed effective properties and suitability for field application trials (Nedveckyte et al., 2021). Moreover, it was found active against *Frankliniella occidentalis* Pergande (Zhou et al., 2020) together with several other *Lecanicillium* spp.

In greenhouse bioassays, *L. aphanocladii* DSF efficacy was measured based on nymphal survival/adult emergence rate of *P. spumarius*, with different combinations of propagules (conidia or blastospores) and adjuvants (adjuvant 1 and 2), compared to both negative control (water) and a commercial product based on *L. muscarium* (Mycotal®) which finally resulted not effective on *P. spumarius* nymphs. All experimental theses with *L. aphanocladii* DSF, except for conidia formulation with adjuvant 1, significantly increased nymphal mortality and reduced adult emergence rate compared to the controls, water and adjuvants alone. Apart for the lower mortality rate caused by *L. aphanocladii* DSF conidia formulation with adjuvant 1, no significant differences between the other *L. aphanocladii* DSF formulations were observed in efficacy against *P. spumarius*. A second trial was carried out to compare the two *L. aphanocladii* DSF formulations (conidia and blastospores) with the commercial product Naturalis® based on a *B. bassiana* strain. Even if all the EPF theses (*L. aphanocladii* DSF and *B. bassiana* Naturalis®) significantly reduced nymphal survival and adult emergence compared to the control, *L. aphanocladii* DSF blastospore formulation was significantly more effective than the conidia formulation, with nymphal mortality/adult emergence rates (mortality: 90%; emergence: 4 %) even higher, although not significantly than those observed in *B. bassiana* theses.

Despite their well-known potential, few authors have tested the efficacy of blastospores, compared with conidia, in pathogenicity bioassays. Blastospores are proposed as suitable contact mycoinsecticide in spray applications for their ability to germinate and infect insects more rapidly than conidia (6h versus 16-24h). At the same time, blastospores are less tolerant to environmental stresses than conidia, even though faster germination may reduce the exposure period to deleterious environmental stresses after application (Jaronsky and Mascarin, 2017). For these reasons, we investigated and compared the pathogenic action of both *L. aphanocladii* DSF propagules. Furthermore, blastopore-based mycoinsecticides of *Lecanicillium* sp. are currently produced (Mycotal® and Vertalec®).

The selection process of a mycoinsecticide must also evaluate the potential of fungal strain to form stable propagules that can be economically mass-produced (Jaronsky and Mascarin, 2017). Stability in storage is critical in most commercial situations, although in some cases provision of fresh mycoinsecticide to the user is feasible. Blastospores are considered less amenable to storage conditions but more sensitive to desiccation than conidia. Nevertheless, cost-effective methods for producing high concentrations of desiccation-tolerant blastospores of *Lecanicillium* and *Beauveria* have been already successfully achieved and reported (Jaronsky and Mascarin, 2017). From our experience, freeze dried blastospores remained stable for months without losing vitality but it was necessary to store them at 4 °C which is inconvenient for industry. Furthermore, the hydrophilic nature of blastospores, compared with hydrophobic conidia, make them extremely difficult to suspend in acqueous carriers, making necessary further efforts in formulation optimization.

In the same trials, we tested also for the influence of spittlebug foam on effectiveness of *B. bassiana* Naturalis® conidia treatment on nymphal stages. No differences in mortality or adult emergence rate were observed between nymphs treated when covered by foam or “naked”, i.e. with no foam. The results, although preliminary, showed that the presence of foam seemed to neither hinder nor favour the pass through of *B. bassiana* hydrophobic conidia. The composition of spittlebug foam is not well-known, but generally resemble the exudate of host plants, although the nymphs secrete several chemical component, such as glycopeptides and proteoglycans, as well as metabolites with likely defensive/repellent function towards predators (Mello et al. 1987; del Campo et al. 2011; Chen et al. 2018). Potential effects of foam composition and structure on effectiveness of EPF should be explored in further studies.

The results obtained in this study represent one of the first evidence on EPF genera and species attacking *P. spumarius*. The initial recovery of *P. spumarius* individuals affected by mycosis was a fundamental step towards a good chance of success. In general, native strains naturally infecting target insects are quite promising in the development of biocontrol agents (Dara et al., 2008; Wang et al., 2022). Indeed, EPF strains isolated from arthropod cadavers show a more extensive natural control ability and a superior environmental tolerance. Some authors highlighted that the virulence of indigenous EPF strains can be higher than that of commercial strains and their effectiveness is largely dependent on genetic variability and adaptation to host related environmental factors (Ou et al., 2019; Wu et al., 2021; Thube et al., 2022). Boucias et al. (2007) started from a field-survey to recover fungi associated to the spittlebug *H. vitripennis*, and finally selected a strain belonging to *H. homalodiscae* as biocontrol agent (Lietze et al. 2011). Foieri et al. (2017) also observed the natural occurrence of genus *Pandora* on spittlebug pests (Hemiptera: Cercopidae) of pasture grasses, although no isolation of fungal pure cultures from individuals affected by mycosis was achieved. The inability (or scarce ability) to grow and sporulate *in vitro* is a major limitation of some EPF, especially belonging to the order of *Entomophthorales*, which makes them unsuitable for possible use. Many species have never been cultured *in vitro* because of their nutritional fastidiousness (Jaronsky and Mascarin, 2017).

Data collected from climatic chamber assays are quite encouraging even if trials were carried out under controlled conditions, optimal for fungus development in terms of temperature and RH. However, we avoided *in vitro* conditions used by other authors (Cabinallas and Jonas 2013; Ganassi et al., 2023) and the experimental set up was closer to environmental conditions. Pathogenicity of EPF strains were tested by direct spraying of fungal propagules on *V. faba* leaves in microcosms colonised by *P. spumarius* to ensure almost complete coverage and maximize the probability of contact and infection of the target insects. The low mortality of nymphs detected in the negative control (Wtr) highlighted a good set up of the trials with good environmental conditions for *P. spumarius* survival.

Certainly, the results obtained in this study require validation in field assays in order to confirm the potential of *L. aphanocladii* DSF against *P. spumarius* nymphs. Environmental conditions are often a key determinant of biopesticide efficacy, as EPF perform best at optimal conditions for both germination and host infection. In addition, due to the relative slowness of EPF in acting, further studies should be focused on the capacity of *L. aphanocladii* DSF in disease transmission (e.g. mycosis-affected individuals stuck to the leaves which can transmit the fungus to healthy individuals) (Lietze et al., 2011) and persistence in the environment (e.g. propagule resistance to UV and fungal aptitude for endophytic colonisation of target insect host plants).

Although the European objectives for eco-sustainability and organic farming are accelerating the introduction of microbial pesticides, their use remains limited and new efforts should be made by the scientific community to test the efficacy of new strains. This study represents a first attempt in answering the question if entomopathogenic fungi may represent an alternative or be synergic to natural insecticides to achieve effective control of spittlebug populations in agroecosystems threatened by *X. fastidiosa*.

## ACKNOWLEDGEMENTS

We thank Liesbeth Zwarts, NV Globachem, Sint Truiden, Belgium, for providing blastospore preparation of *Lecanicillium aphanocladii*. We wish to thank Matteo Alessandro Saladini, Emanuela Di Vasto and Clara Ontañon for technical support in mortality assays.

