## Supplemental Table 1 for "Activity of natural occurring entomopathogenic fungi on nymphal and adult stage of *Philaenus spumarius*"

### SUPPLEMENTARY MATERIAL

*Table S1: T2-2022; Hazard ratios and 95% confidence intervals (95% CI) for transitioning of Philaenus spumarius nymphs from alive to dead and alive to adult emergence for experimental theses (EPF treatments + control)..*

| **Transition** | **Treatment** | **With respect to** | **Hazard ratio** | **95% CI** | | **z** | **p-value** |  |
| --- | --- | --- | --- | --- | --- | --- | --- | --- |
| Alive to dead | *LaB* | *Wtr* | 16.56 | 7.07 | 38.76 | 6.59 | <0.001 | *** |
|  | *LaC* | *Wtr* | 4.93 | 2.13 | 11.44 | 3.7 | <0.001 | *** |
|  | *NCf* | *Wtr* | 11.14 | 4.86 | 25.54 | 6.05 | <0.001 | *** |
|  | *NCbf* | *Wtr* | 14.13 | 6.17 | 32.35 | 6.82 | <0.001 | *** |
|  | *Wtr* | *LaB* | 0.08 | 0.04 | 0.17 | -6.59 | <0.001 | *** |
|  | *NCbf* | *LaB* | 0.98 | 0.65 | 1.5 | -0.07 | 0.94 |  |
|  | *NCf* | *LaB* | 0.75 | 0.48 | 1.18 | -1.24 | 0.21 |  |
|  | *LaC* | *LaB* | 0.33 | 0.2 | 0.53 | -4.53 | <0.001 | *** |
|  | *LaC* | *NCf* | 0.43 | 0.28 | 0.67 | -3.71 | <0.001 | *** |
|  | *LaB* | *NCf* | 1.33 | 0.85 | 2.07 | 1.24 | 0.21 |  |
|  | *Wtr* | *NCf* | 0.11 | 0.05 | 0.22 | -6.05 | <0.001 | *** |
|  | *NCbf* | *NCf* | 1.3 | 0.91 | 1.88 | 1.44 | 0.15 |  |
|  | *LaB* | *LaC* | 3.07 | 1.89 | 4.98 | 4.53 | <0.001 | *** |
|  | *Wtr* | *LaC* | 0.25 | 0.12 | 0.52 | -3.7 | <0.001 | *** |
|  | *NCbf* | *LaC* | 3.02 | 1.97 | 4.63 | 5.08 | <0.001 | *** |
|  | *NCf* | *LaC* | 2.31 | 1.49 | 3.61 | 3.71 | <0.001 | *** |
|  | *NCf* | *NCbf* | 0.77 | 0.53 | 1.1 | -1.44 | 0.15 |  |
|  | *LaC* | *NCbf* | 0.33 | 0.22 | 0.51 | -5.08 | <0.001 | *** |
|  | *LaB* | *NCbf* | 1.02 | 0.67 | 1.55 | 0.07 | 0.94 |  |
|  | *Wtr* | *NCbf* | 0.08 | 0.04 | 0.17 | -6.82 | <0.001 | *** |
| Alive to adult | *LaB* | *Wtr* | 0.16 | 0.03 | 0.69 | -2.067 | 0.039 | * |
|  | *LaC* | *Wtr* | 0.52 | 0.29 | 0.93 | -2.132 | 0.033 | * |
|  | *NCf* | *Wtr* | 0.47 | 0.21 | 1.05 | -2.77 | 0.006 | ** |
|  | *NCbf* | *Wtr* | 0.37 | 0.13 | 1.05 | -2.927 | 0.003 | ** |
|  | *Wtr* | *LaB* | 10.02 | 2.14 | 46.85 | 2.93 | <0.001 | *** |
|  | *NCbf* | *LaB* | 3.33 | 0.52 | 21.44 | 1.27 | 0.2 |  |
|  | *NCf* | *LaB* | 4.22 | 0.77 | 23.09 | 1.66 | 0.1 |  |
|  | *LaC* | *LaB* | 4.52 | 0.94 | 21.76 | 1.88 | 0.06 |  |
|  | *LaC* | *NCf* | 1.07 | 0.45 | 2.55 | 0.15 | 0.88 |  |
|  | *LaB* | *NCf* | 0.24 | 0.04 | 1.29 | -1.66 | 0.1 |  |
|  | *Wtr* | *NCf* | 2.37 | 1.07 | 5.24 | 2.13 | 0.03 | * |
|  | *NCbf* | *NCf* | 0.79 | 0.25 | 2.52 | -0.4 | 0.69 |  |
|  | *LaB* | *LaC* | 0.22 | 0.05 | 1.07 | -1.88 | 0.06 |  |
|  | *Wtr* | *LaC* | 2.22 | 1.26 | 3.9 | 2.77 | 0.01 | * |
|  | *NCbf* | *LaC* | 0.74 | 0.24 | 2.23 | -0.54 | 0.59 |  |
|  | *NCf* | *LaC* | 0.94 | 0.39 | 2.23 | -0.15 | 0.88 |  |
|  | *NCf* | *NCbf* | 1.27 | 0.4 | 4.04 | 0.4 | 0.69 |  |
|  | *LaC* | *NCbf* | 1.35 | 0.45 | 4.09 | 0.54 | 0.59 |  |
|  | *LaB* | *NCbf* | 0.3 | 0.05 | 1.93 | -1.27 | 0.2 |  |
|  | *Wtr* | *NCbf* | 3 | 1.06 | 8.52 | 2.07 | 0.04 | * |
